# DNA metabarcoding for high-throughput monitoring of estuarine macrobenthic communities

**DOI:** 10.1101/117168

**Authors:** Jorge Lobo, Shadi Shokralla, Maria Helena Costa, Mehrdad Hajibabaei, Filipe Oliveira Costa

## Abstract

Benthic communities are key components of aquatic ecosystems’ biomonitoring. However, morphology-based species identifications remain a low-throughput, and sometimes ambiguous, approach. Despite metabarcoding methodologies have been applied for above-species taxa inventories in marine meiofaunal communities, a comprehensive approach providing species-level identifications for estuarine macrobenthic communities is still lacking. Here we report a combination of experimental and field studies demonstrating the aptitude of COI metabarcoding to provide robust species-level identifications within a framework of high-throughput monitoring of estuarine macrobenthic communities. To investigate the ability to recover DNA barcodes from all species present in a bulk DNA extract, we assembled 3 phylogenetically diverse communities, using 4 different primer pairs to generate PCR products of the COI barcode region. Between 78 and 83% of the species in the tested communities were recovered through HTS. Subsequently, we compared morphology and metabarcoding-based approaches to determine the species composition from four distinct sites of an estuary. Our results indicate that the species richness would be considerably underestimated if only morphological methods were used. Although further refinement is required for improving the efficiency and output of this approach, here we show the great aptitude of COI metabarcoding to provide high quality and auditable species identifications in macrobenthos monitoring.

## Introduction

Macrobenthic invertebrate surveys have been widely used for the assessment of the ecological status of aquatic ecosystems worldwide^1,2,3,4,5^. They are one of the key compulsory components of biological monitoring programs implemented in numerous countries’ environmental directives, such as the European Union Water and Marine Strategy framework directives (WFD 200/60/EC and MSFD 2008/56/EC) or the USA (EPA 841-B-99-002) and Canadian Aquatic Biomonitoring Network^6^. Under the WFD, for example, EU member states are required to establish a regular biological monitoring programme for freshwater systems and transitional waters (e.g. estuaries), which include macrobenthic communities^7^. So far, routine assessments of macrobenthic invertebrates have been carried out using almost exclusively morphology-based approaches for species identifications. This is time-consuming and skill-dependent approach, which has resulted in low-throughput in processing biomonitoring samples. Very often the specimens cannot be accurately assigned to species, either because morphology-based identifications are inherently difficult, or because of organisms’ bodies that are damaged and missing diagnostic parts. In the case of immature stages, the species level identifications are extremely difficult or nearly impossible^8^. Published data on macrobenthic surveys frequently report specimens assigned only to family or genus level^9,10^. Moreover, in most studies the reliability of the species level identifications cannot be ascertained because the specimens are discarded. The growing reports of cryptic species among dominant macrobenthic invertebrates further calls into question the precision of morphology-based identifications^11,12,13^. However, many of the biotic indexes applied to macrobenthic communities require species-level identifications, such as for example the Azti Marine Biotic Index (AMBI), which is based on a list of nearly 8000 species that are assigned to five ecological groups depending on each species’ tolerance to environmental disturbance^14^.

Recent DNA-based approaches to species identifications from bulk community samples, such as environmental DNA barcoding or DNA metabarcoding^15,16^, have the potential to help circumvent many of the above-described limitations of the morphology-based method. DNA metabarcoding is expected to help improving macrobenthic surveys, by providing a high-throughput approach that generates auditable species-level identifications. Although proof of concept studies have shown the feasibility of the application of metabarcoding approaches for monitoring river macrobenthos^15,17^, no equivalent comprehensive studies have been developed specifically for marine and estuarine macrobenthic communities. Most of the published HTS based studies in estuarine ecosystems targeted genomic regions where species level resolution is limited^18^ and focused exclusively on meiofaunal communities, which used environmental DNA (eDNA) from the sediment^19,20,21^ rather than bulk communities. So far, only a few studies have applied DNA metabarcoding to marine macrobenthic communities using the standard cytochrome c oxidase I (COI) barcode region^22,23,24^. Yet, these few studies either did not test the amplification success of different primer pairs in engineered communities of known species composition for methodology validation, and/or or did not target specifically estuarine soft-bottom macrobenthos. Due to their high phylogenetic diversity, marine and estuarine communities may convey additional difficulties in PCR-based approaches due to potential primer mismatch and amplification bias^25,26,27^, therefore specific approaches must be comprehensively tested before conducting full “blind” metabarcoding assessments^22^.

Our aim in the present study is a) assess the ability of metabarcoding approach to detect the diversity of species typically present in estuarine macrobenthic communities, through the use of experimentally assembled communities; b) evaluate whether the metabarcoding approach provides comparable, more or less detailed species inventories compared to the traditional morphology-based approaches; and c) assess the ability of the metabarcoding approach to effectively discriminate among natural macrobenthic communities within an estuary, therefore reflecting environmental conditions at different sites and enabling its use in the assessment of the sediment environmental quality and ecological status of the estuary. By combining experimental and field studies, here we demonstrate for the first time the feasibility of using COI metabarcoding for monitoring estuarine macrobenthic communities, which provides equal or more sensitive data on the species composition compared to morphology.

## Results

### Metabarcoding of the assembled communities

Species detection success through metabarcoding of the assembled microbenthic communities (AMC) was generally high, ranging from 78% of the species in AMC1 to 83% in AMC3, with 81% of the species detected in AMC2. Two non-target species were detected in the AMC1 although they were included in other AMC - *Abra alba* (W. Wood, 1802) in AMC2 and AMC3 and *Scolelepis (Scolelepis) foliosa* (Audouin & Milne Edwards, 1833) included in AMC2 (see Table 1). The cumulative success of recovery, considering all species present in the three AMC, was 83%; it increases to 89% if we consider the non-target species. Primer pairs were not equally effective for species detection in all three AMC. Globally, the most effective primer was D (78%), followed by B (75%), C (61%) and A (44%) (Fig. 1). Primer pairs B and D showed significant differences with the primers A and C. Notably, the global maximum success rate of species detection was attainable using only two primer pairs, B and D. Six species were not detected with any primer pair, namely three crustaceans (*Corophium* sp.3, *Cyathura carinata* (Krøyer, 1847) and *Melita palmata* (Montagu, 1804)) and one polychaete (*Scolelepis sp*.). One bivalve (*Abra alba*) and one polychaete species (*Scolelepis (Scolelepis) foliosa*) have to be added if the non-target species are considered.

**Figure 1:**
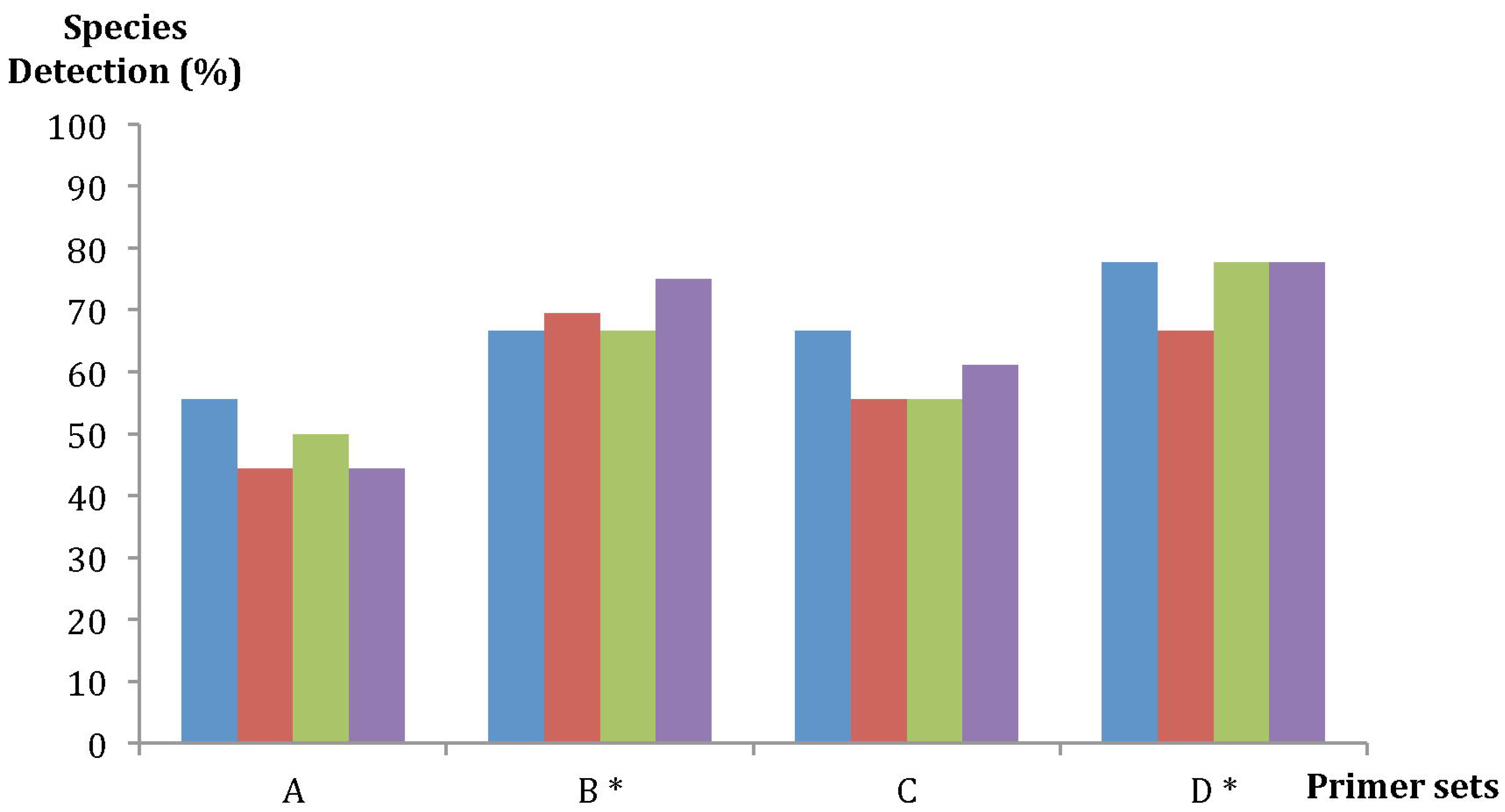
Species detection success for the four primer pairs (A, B, C and D). The columns in each primer pair (from left to right) denote: AMC1, AMC2, AMC3 and global result for the three AMC.

**Table 1.**
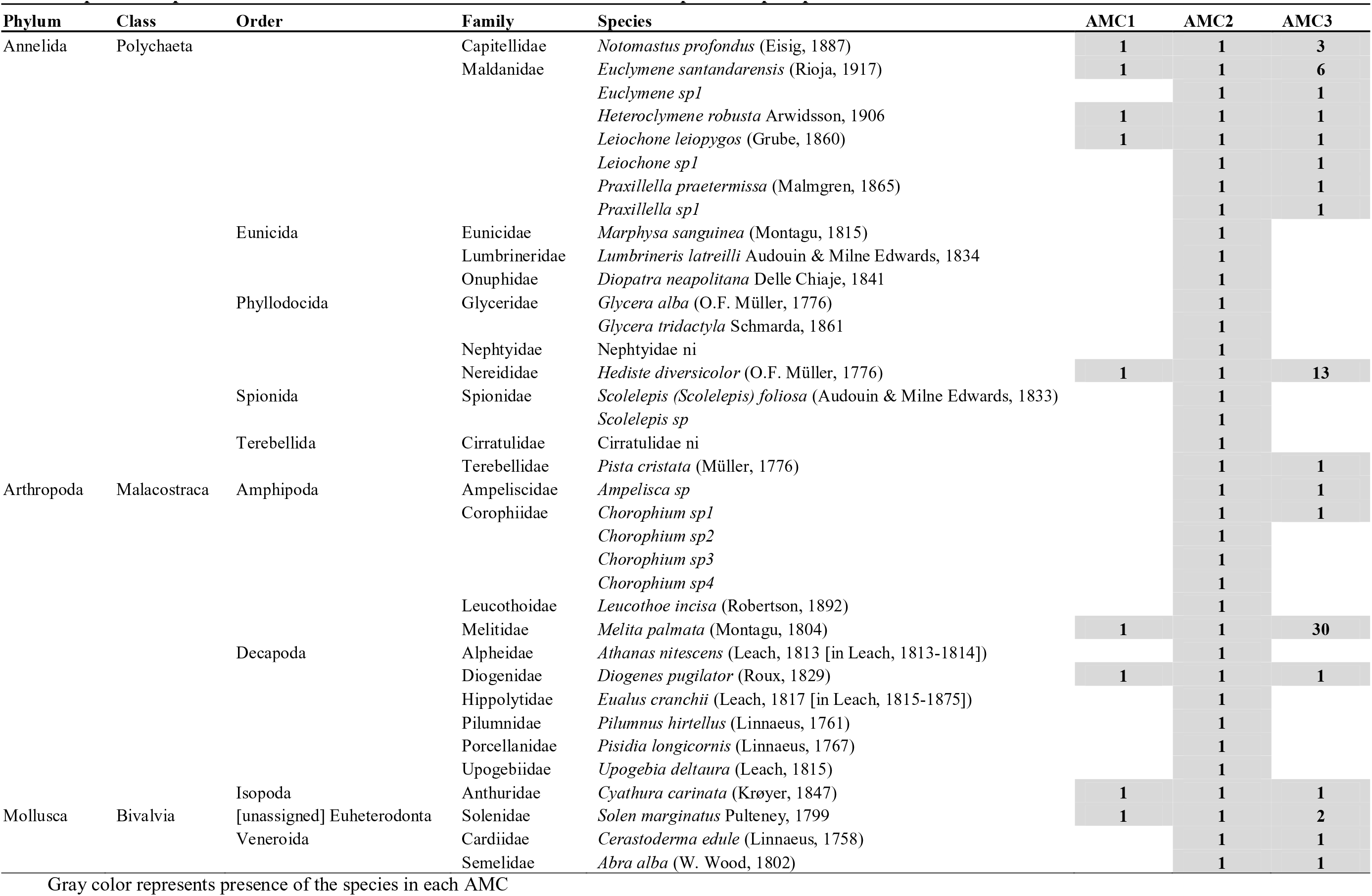
Species composition of the three assembled communities and number of specimens per species

### Metabarcoding of the natural communities

The sediments’ types in the 4 sites of the Sado estuary sampled for the natural macrobenthic communities (NMC) varied considerably in their features, ranging from sandy to muddy sediments (Table 2). The sediment at the NMC1 and NMC3 sites had respectively the lowest and highest TOM content among the four sediments analyzed. Sediments of the NMC2 and NMC3 also had high organic matter content (1.30% and 2.05%, respectively), however, NMC2 had lower FF because was probably disturbed due to the recent construction of the ferryboat wharf. The results are summarized in Table 2.

**Table 2.** Sediment features in the 4 sites of the Sado estuary sampled for the natural macrobenthic communities

|  | NMC1 | NMC2 | NMC3 | NMC4 |
| --- | --- | --- | --- | --- |
| **Salinity** | 34 ± 1 | 34 ± 1 | 34 ± 1 | 34 ± 1 |
| **TOM (%)** | 0.62 ± 0.05 | 1.30 ± 0.11 | 2.05 ± 0.20 | 0.74 ± 0.18 |
| **FF<sup>a</sup> (%)** | 5.16 | 6.4 | 16.93 | 9 |

TOM total organic matter, FF fine fraction
<sup>a</sup> Particle size < 63 µm

Morphological identification of the specimens was carried out in five corer samples per site, except for NMC2, where no specimens were found after sieving one corer (NMC2.10). Species level identifications were attempted in the majority of specimens. except those taxonomically difficult groups (e.g. oligochaetes and nemerteans) and the very damaged or fragmented specimens due to the sieving process. A few specimens of polychaetes (family Cirratulidae and genera *Euclymene* and *Glycera*) and amphipods (genus *Ampelisca*) were classified to higher taxonomic ranks, since these taxa are especially difficult to identify through traditional methods. Considering only the specimens identified to the species level, four phyla were detected in all NMC (Annelida, Arthropoda, Echinodermata and Mollusca), but this number increased to five (plus Nemertea) if we consider specimens identified to a higher taxonomic level. All communities showed a diverse taxonomic composition, comprising between 3 and 5 phyla, except NMC1, which was only composed of polychaetes and mollusks (Supplementary Fig. S1A, B). Globally, 55 taxa were identified in the natural communities, 27 of which were identified to species level and the remaining 28 to higher taxonomic ranks.

Metabarcoding-based identification generated a total of 61 species matches in all 4 natural communities, obtained through searches against both GenBank public database and our own reference library (dx.doi.org/10.5883/DS-3150). The 61 species were distributed among six phyla, the same 5 reported above from the morphological identification, plus Bryozoa. The variation of the species richness among sites displayed a similar pattern in morphology or metabarcoding-based assessments (NMC2 < NMC1 < NMC3 = NMC4 for morphological identifications and NMC1 < NMC2 < NMC3 = NMC4 for metabarcoding) but the number of species recorded was more than twice using the latter method (Supplementary Fig. S1C). NMC1 was also the less taxonomically diverse together with NMC2, represented only by three phyla. Forty-three of the 61 species were detected by any of the primer pairs used (B and D). Among the remaining 18 species, 10 were recovered exclusively with primers B and 8 exclusively with the primers D. The number of reads assigned to species in each sample of all NMC and primer pair is available as Supplementary Table S1.

Comparison between morphological and metabarcoding species-level identifications in the 4 natural macrobenthic communities resulted in only 23% (range 20-28%) of the species detected simultaneously by the 2 approaches (Fig. 2). In average, as much as 65% of the species were detected exclusively by metabarcoding (range 62%-71%), while species detected exclusively by morphology were only 12% in average (range 9%-15%). Among the latter, there were 4 species for which there were no reference COI barcodes available when the analysis was performed (*Corbula gibba* (Olivi, 1792), *Ecrobia ventrosa* (Montagu, 1803), *Spisula solida* (Linnaeus, 1758) and *Parvicardium pinnulatum* (Conrad, 1831)). Polychaetes were the dominant taxa in all sites, regardless identified by morphology or metabarcoding, except for morphology based identifications in NMC2, that were dominated by molluscs. The second most important groups were arthropods in the case of metabarcoding-based identifications, and molluscs in the case of morphology-based identifications. The detailed list of species identified in each site by the two approaches is available as Supplementary Table S2, while the proportion of taxa in each corer and approach is displayed in Supplementary Fig. S1.

**Figure 2:**
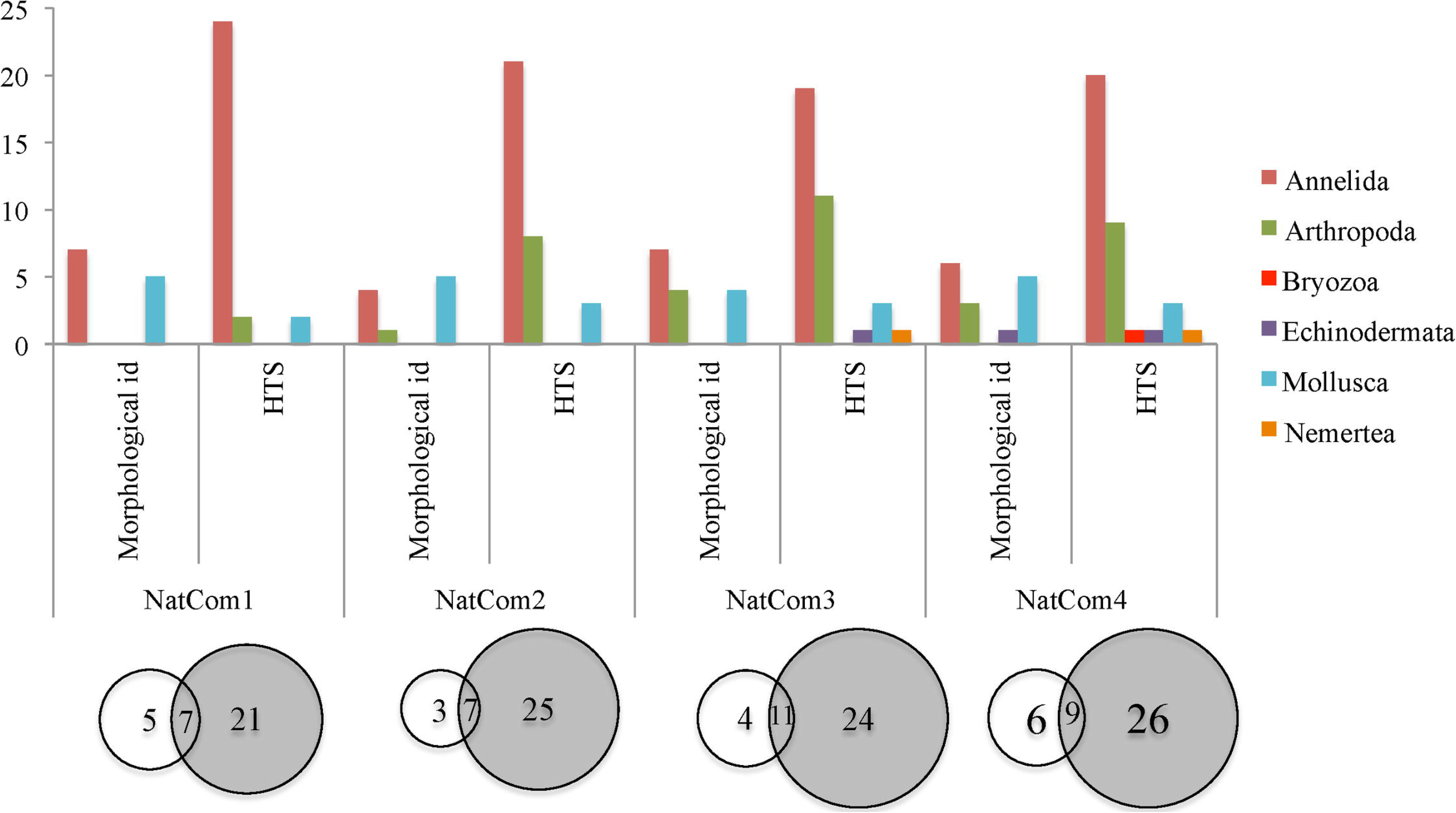
Comparison between morphological and metabarcoding species-level identifications in 4 macrobenthic communities (NMC1-NMC4) of the Sado estuary, Portugal. The upper bar chart shows the distribution of the number of species per phylum obtained either by morphology or metabarcoding, in each macrobenthic community. The circles in the lower part of the figure represent the proportion of species detected exclusively by morphology (white circles), exclusively by metabarcoding (shaded circles), and by both approaches (overlapping circles dashed area).

Fig. 3 shows the graphical distribution of the natural communities as a function of their similarities in species composition, obtained by non-parametric MDS, for either the morphology, metabarcoding-based identifications and also the combination of two approaches. Three maps show a similar pattern, NMC1 and NMC2 in the left part of the map and NMC3 and NMC4 in the right side. The combination of the two identification approaches approximates even more the NMC1 and NMC2. The results obtained using AMBI also showed a similar pattern between the morphological identification, HTS and combination of both approaches, all calculated using only absence-presence of species. On the other hand, the original AMBI index also showed similar results with the AMBI index using absence-presence of species for the morphology-based identifications. The four NMC were classified as slightly disturbed probably because the majority of the species obtained in each natural community through the three approaches was similar. Although NMC1 was the community closer to the EG-III (moderately disturbed) and NMC2 and NMC3 the less disturbed (see Fig. 4).

**Figure 3:**
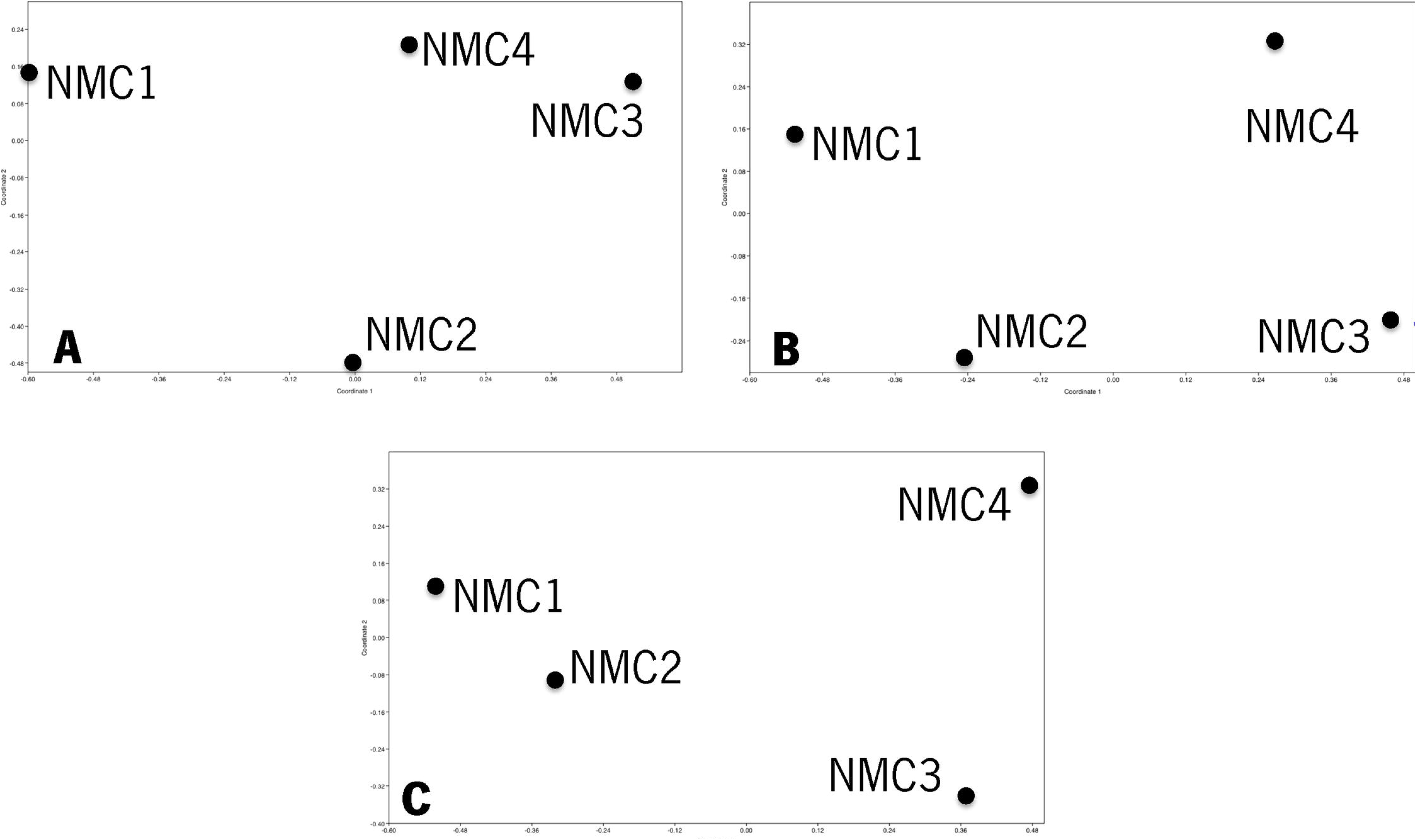
Non-metric multidimensional scaling (nMDS) for the morphological identification (A), HTS (B) and morphological identification plus HTS (C) results of the four NMC. Similarity index of Bray-Curtis was applied for the absence-presence of the species.

**Figure 4:**
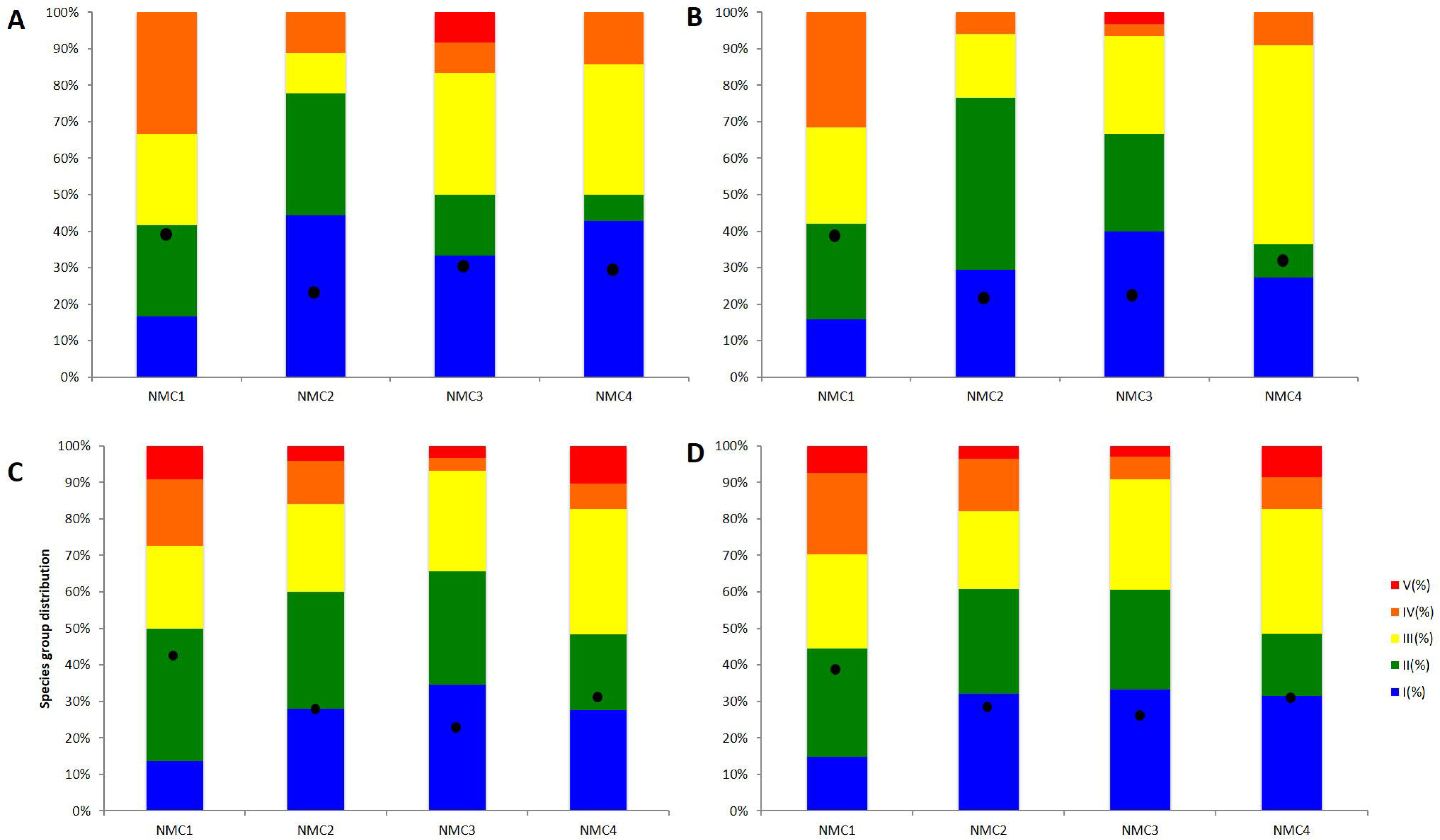
Comparison of AMBI for the morphological identification using absence-presence of species (A), morphological identification using abundance of species (B), HTS (C) and morphological identification plus HTS (D) results of the four NMC.

## Discussion

The combination of samples representing assembled communities and field collected bulk samples demonstrates the potential for implementing COI metabarcoding in the monitoring of estuarine macrobenthic communities. The tests performed with assembled communities of known composition, showed that high success rates in species detection are attainable using COI amplicons and employing only two primer pairs. In the field tests, COI metabarcoding generated concordant results with morphology based assessments, and detected a higher number of species in all stations and samples. Finally, the metabarcoding approach was sensitive and able to reflect differences in the species composition among natural communities.

In spite of the differences in the proportion of specimens per species, relatively high success rates in species detection were attained in all of the assembled communities (78% to 83%). AMC2, composed of the highest number of species (36), each represented by a single specimen, constituted an extreme test for the robustness of the metabarcoding approach, particularly for the effectiveness of the bulk DNA extraction and amplification procedures. In this community, no sequences were generated only for two species (*Corophium* sp. 3 and *Scolelepis* sp.) with any of the 4 primer pairs tested. Specimens of these species were very small (< 5mm in length) and the possibility that their DNA was not effectively isolated and that not enough template DNA was available for amplification cannot be discarded. Two species of peracaridean crustaceans (*Cyathura carinata* and *Melita palmata*) were apparently recalcitrant to amplification generating no reads, although they were present in the three assembled communities. However, the fact that previously we have been able to generate full DNA barcodes for individual specimens by Sanger sequencing using one of the primer pairs (Lobo primers^25^), and that the isopod *C. carinata* was recovered in the natural communities, excludes the possibility of amplification inhibition in these species. A possible explanation is that these species have a low affinity to the tested primer pair and may be outcompeted by higher affinity DNA templates from other species present in the PCR reaction. This is an important issue when considering primer match for metabarcoding studies and demonstrates the need for primer evaluation using assembled mixtures prior to large-scale analysis of bulk samples. Several reasons could explain the detection of two species (*Abra alba* and *Scolelepis (Scolelepis) foliosa*) in AMC1 where they were not included, but not in AMC2 and AMC3, where they present. Because the organisms were processed in the same collection event, some tissue or body fragment may have been accidentally transported together with other specimens, or they may have even been preyed upon by some of the predator species (e.g. *Hediste diversicolor*) present in AMC1.

Mismatches between primers and target templates are a key concern in PCR-based metabarcoding, since it can lead to some level of systematic failure in species detection^28^. Because in silico analyses reveal high variability in the actual and potential primer annealing regions within the COI barcode, this marker has been dismissed as appropriate for metabarcoding^29^. Alternative markers, such as the nuclear gene coding for 18s rRNA, with lower variability in priming sites, have been used and proposed for metabarcoding marine macroinvertebrates^22,30^, but the species level resolution is substantially lower than using COI^19,22,26^. Additionally, 18s rRNA primers are not free of PCR-bias. When compared side by side in a field test of metabarcoding invertebrates of seagrass meadows^22^, both markers showed taxonomic bias, with the 18s rRNA recovering a higher number of species (compared to full length COI barcodes (658 bp)) but amplifying preferentially meiofaunal groups such as nematods. Since species level identification is essential for applying macrobenthic invertebrate indices (e.g. AMBI^14^), and reference libraries for marine invertebrates are available and continuously growing, the standard barcode marker for metazoans is the natural candidate for metabarcoding macroinvertebrate communities. Several studies^31^ have shown that shortcomings of PCR bias may be minimized by using enhanced degenerate primers, and multiple amplification primers. The deep sequencing provided by the HTS platforms used in metabarcoding may also improve global primer success compared to what has been found using individual specimen sanger sequencing^31,32,33^.

Despite no major differences were observed in species detection success rates among the three different assembled communities, there were considerable differences among the 4 primer pairs. The primer pair A, amplifying 658 bp, was the least successful one; hence we conclude that smaller length sizes appear to be more efficient for metabarcoding. Short fragments of COI barcode (mini-barcodes), even of 150 bp, can achieve unambiguous species-level identifications, as it was observed for a diversity of taxonomic assemblages in previous studies^32,33,34,35^. A much better success rate was obtained with primer pairs B and D compared to A and C. The two former primers combinations were here tested for the first time, and proved effective in the amplification of more target species from three phyla than the remaining two primers. There was also no indication of a major taxonomic bias in these primers, as they were able to amplify targets from any of the three phyla. This indicates that, despite the large phylogenetic diversity of estuarine communities, a combined approach of degenerate primer design and multiple amplification primers can minimize substantially primer-template mismatch issues.

No relationship was found in this study between the number of specimens and the number of reads. Indeed, for phylogenetically diverse assemblages such as macrobenthic communities, comprising organisms varying widely in size, biomass and anatomically (thus varying also in the amount of DNA template available in a bulk extraction), the possibility of quantitative inferences from the number of reads data was not anticipated. For example, the polychaete species *Hediste diversicolor*, represented by 1 specimen in the AMC1 and 14 specimens in the AMC3, produced 8 and 23194 reads respectively, using the primer pair B. However, the polychaete species *Notomastus profondus*, represented by 1 specimen in the AMC1 and 3 specimens in the AMC3, produced 4601 and 3161 reads respectively, using the primer pair D. Also, in the AMC2 where all species were represented by 1 specimen, 6131 and 21576 reads of the similar-sized decapod specimens of *Pilumnus hirtellus* and *Upogebia deltaura* were respectively obtained, among various other examples of deep mismatches between the number of reads and organisms abundance and size patent in our results. Empirical relationships between specimen numbers, body size or biomass and the number of reads have been found occasionally, usually in studies targeting a closely related and more or less homogeneous group (e.g. chironomids^36,37^). In comprehensive tests performed by Elbrecht and Leese^38^ such relationships were still elusive, probably because the primer efficiency is highly species-specific, preventing straightforward inference of species abundance in the assembled communities.

Marine macrobenthic communities are complex, highly diverse communities, where morphology-based species identifications can be rather challenging. In our study many specimens could not be identified to the species level due to uncertain species identity, mostly when they were immature stages, belonged to difficult taxa or were missing diagnostic body parts as a result of sieving, handling and ethanol. Such difficulties are common, even when a group of experts is available, as reported in numerous studies^39^. We have found many fragments of organisms, namely of annelids, as a result of sieving and handling process and therefore many species could not have been identified using morphology, although they were present in the samples. A comprehensive search over 138 published reports and inventories of benthic communities has found that approximately one third of the specimens were not identified to species level when using morphological methods, although this proportion of missed species identifications varies greatly between different taxonomic groups^40^. The morphology-based macrobenthic community profile that we obtained for the four sites in the Sado estuary, provided similar results to previous morphology-based surveys in nearby and similar ecosystems, both regarding the species richness and species-specific composition (e.g. Tagus estuary^41^).

In our study, metabarcoding approaches for identifying species composition in communities indicated that the species richness would be underestimated if we used only morphological methods. Similar findings have been reported in a study made on seagrass associated benthic communities^22^, where HTS-based species inventories were considerably richer compared to morphology-based ones. The advantage of using DNA barcodes for metabarcoding approaches is that the reference libraries are being established and improved for all major groups of eukaryotic organisms. Thereby, it is possible to verify the species attribution of the samples. Contrary to the morphological approach, HTS allowed to recover sequences from damaged specimens, immature stages, difficult taxonomic groups, fragments of organisms and even from endoparasites, namely the copepod *Mytilicola orientalis* Mori, 1935 and the decapod *Pinnotheres pisum* (Linnaeus, 1767). *M. orientalis*, native to East Asia, occurs in the intestinal tracts of bivalve species and has been recorded as an alien species in European waters^42,43^. Metabarcoding could be used as a tool for early detection of invasive species^44^. *P. pisum*, living in the mantle cavity of bivalves, is also a parasite^45^. In addiction, six species of algae were recovered: *Pyropia haitanensis* (T.J.Chang & B.F.Zheng) N.Kikuchi & M.Miyata, 2011, *Ceramium secundatum* Lyngbye, 1819, *Durvillaea* sp., *Leathesia marina* (Lyngbye) Decaisne, 1842, *Petalonia fascia* (O.F.Müller) Kuntze, 1898 and *Scytosiphon lomentaria* (Lyngbye) Link, 1833 (see Supplementary Table S2). Although the algae were not a targeted taxonomic group, this illustrates that studies with different scopes are possible, even when using the primer pairs applied in this work.

As presented in Fig. 4, the four natural communities presented each their own species composition. NMC1 and NMC2 appeared to be more similar to each other (on the left side of the graphic) and the same for NMC3 and NMC4 (right side of the graphic), agreeing with their geographic vicinity, on either the north or south margin of the estuary. The species richness was consistently higher in NMC3 and NMC4 for both morphological and HTS approaches. According to the original AMBI and p/a AMBI indexes, the four NMC were classified as slightly disturbed. Sado estuary is globally considered a slightly disturbed ecosystem due to its high hydrodynamics and multiple anthropogenic activities, although the south margin is less disturbed than the north one^46^. Contrary to these global patterns, our AMBI and species richness results indicate NMC1, located in the south margin, as the most disturbed community in this study. Regular dredging operations and construction works, together with a strong hydrodynamics, can affect and promote sudden changes in macrobenthic communities in the Sado estuary^47,48^ and may help to explain these results. However, the key finding is that either morphological or metabarcoding approaches produced similar global outcomes (AMBI classifications and species richness ranks), and metabarcoding consistently outperformed morphology in the ability to detect a higher number of species and to provide species level identifications, despite the still incipient state of completion of the reference libraries for macrobenthic invertebrates.

In summary, our study demonstrates the aptitude of COI metabarcoding using HTS approach for implementation in biodiversity assessments of estuarine macrobenthic communities. High-throughput metabarcoding may enable more frequent and spatially detailed biomonitoring with higher information content^6,17^, concomitantly reducing time and cost constraints in the monitoring of benthic communities. By virtue of the generation of readily comparable DNA sequence data, the metabarcoding approach can provide species-level information of high quality, with reduced ambiguity and susceptible to scrutiny in the future. The ability to provide data on parasite occurrence, for example, and to enable early detection of alien species, or to discriminate cryptic species, constitute highly relevant additional benefits of this approach. Nevertheless, further refinement is still required, to improve its overall efficiency and output, namely the improvement of the recovery rates through the refinement of primers and testing of alternative combinations, especially for the recalcitrant species. Given that the direct measurement of species abundance is still not attainable, further studies are required to generate large datasets, which will allow extensive comparison of the performance of morphology and metabarcoding-based approaches._Lastly, the continuing completion of the still incipient reference libraries of DNA barcodes for marine invertebrates will be decisive to fully materialize the potential of metabarcoding.

## Methods

### Ethics statement

The areas sampled in the Sado estuary do not have any protection status and therefore do not require authorization to carry out scientific work, such as sediment sampling. The sampled macrobenthic fauna does not include any protected or endangered species.

### Experimental design

This study was designed in two main sequential phases. The first phase focused on analysis of macrobenthic communities with known composition, while the second phase comprised natural field-collected macrobenthic communities. In the first phase, the ability of four combinations of primer pairs to successfully amplify fragments of the COI barcode region between 250 to 658 base pairs (bp), was tested in three assembled microbenthic communities. The assembled communities included a different number of species and individuals per species. The two most efficient primer pairs were then used in the second phase. A schematic overview is presented in Fig. 5.

**Figure 5:**
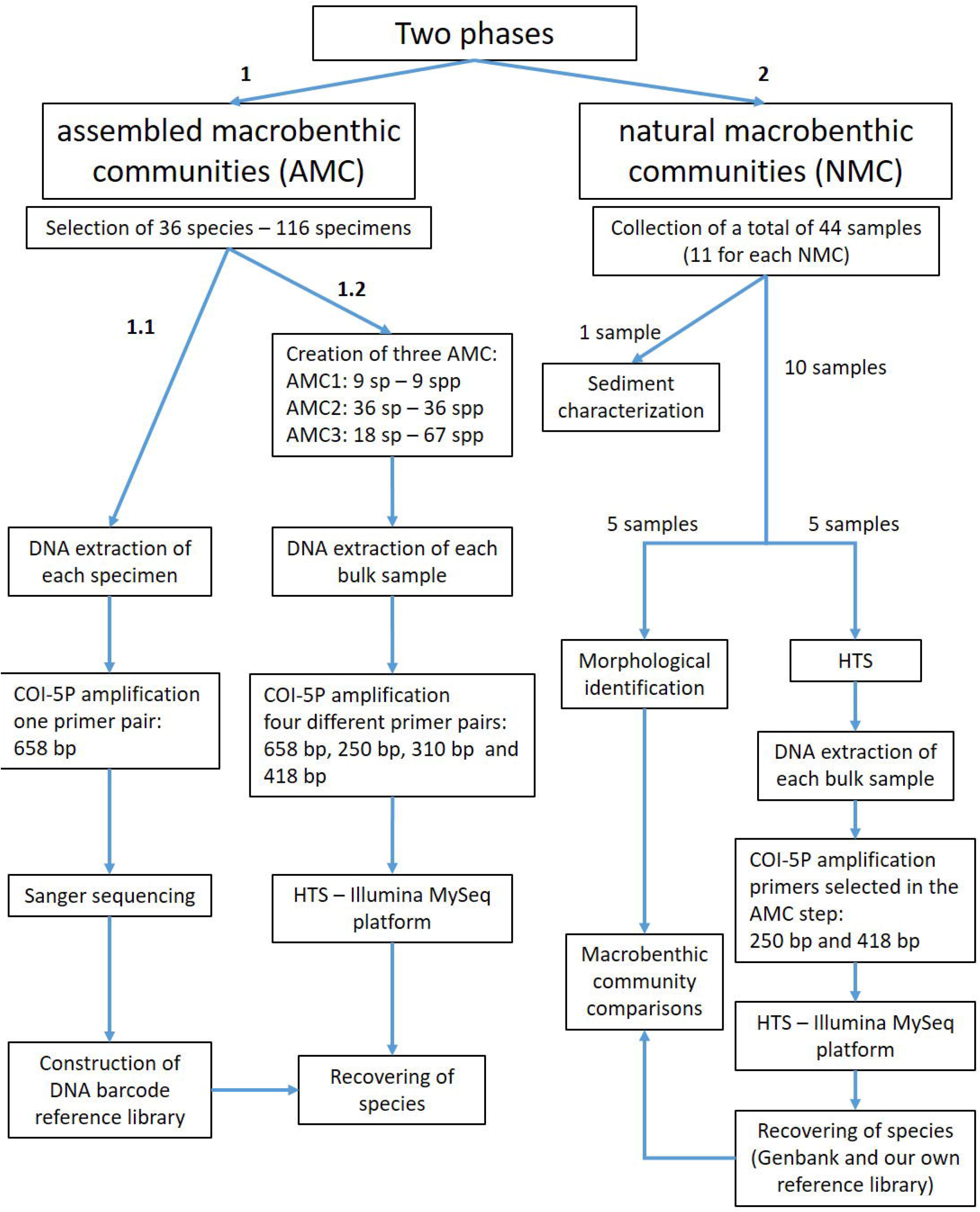
Schematic overview of the experimental design.

In the second phase, morphology-based taxonomic identification of the species composition in the natural macrobenthic communities was directly compared with the species inventory obtained from HTS of COI amplicons generated from bulk community DNA extractions, using two primer pairs selected among the four previously tested. This comparison was applied to NMCs collected in four separate sites in the Sado estuary, Portugal, encompassing distinct sediment features and levels of anthropogenic impact (Fig. 6). In each site, half of the replicate samples were used for conventional morphology-based identification while the remaining half was used for metabarcoding community analyses. Because data generated through metabarcoding does not provide a direct measure of specimen abundance, we used a biotic index based solely on the presence and absence of species to compare morphology and metabarcoding approaches.

**Figure 6:**
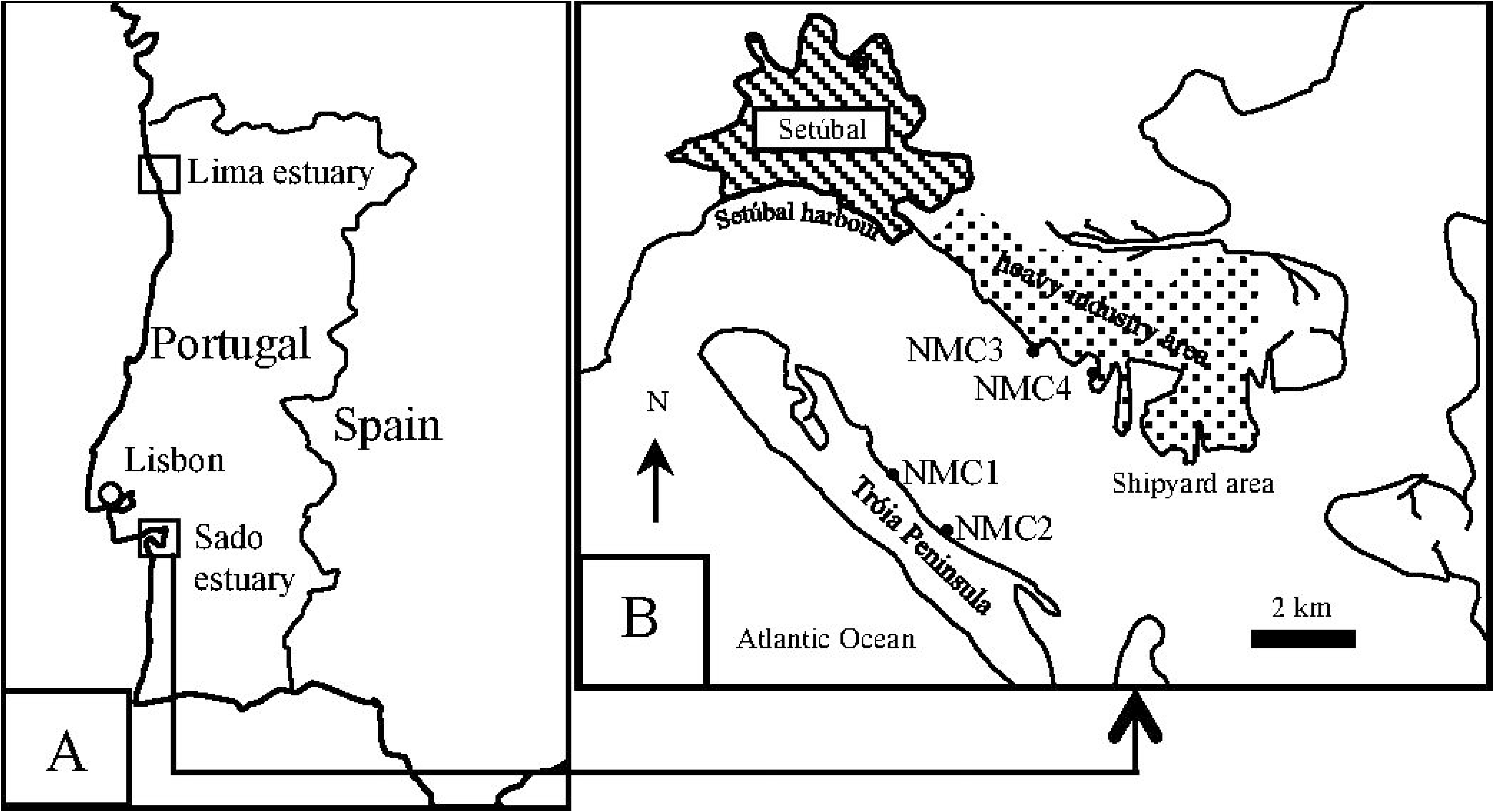
Map of the study area showing the collection sites. A) for the creation of artificial communities and B) for natural communities. NMC = natural communities.

To enable species-level DNA based identifications from the NMC, a reference library of DNA barcodes was compiled for dominant groups of Atlantic European macrobenthic invertebrates. The reference library (dx.doi.org/10.5883/DS-3150) comprises GenBank-published13,25,49,50,51 and unpublished DNA barcodes of marine invertebrates of southern European Atlantic coast, plus the DNA barcodes generated for the specimens used in the AMC study.

### Sediment and specimen collection

#### Assembled macrobenthic communities (AMC)

Specimens were collected in the Sado (Geographical coordinates 38.49/−8.84) and Lima (Geographical coordinates 41.68/−8.82) estuaries, west coast of Portugal (Fig. 6A) during April, May, September and October 2012. Sediment samples were collected using a corer sampler (110 mm diameter, 495 mm height) and sieved through a 0.5 mm screen in order to separate the macrobenthic invertebrates. Sieved samples were transported refrigerated to the laboratory where they were individually separated from the debris and stored in absolute ethanol at −20°C until processing. Morphological identifications to the lowest possible taxonomic level were carried out employing a stereomicroscope, using identification keys^52,53,54,55^. Species’ names were checked in the online databases World Register of Marine Species (http://www.marinespecies.org) and Integrated Taxonomic Information System (www.itis.gov). A total of 112 specimens belonging to 36 morphospecies (25 of which identified to species level) were assembled, comprising 3 mollusks, 13 crustaceans and 20 annelids species, therefore representing the 3 most dominant taxa in typical estuarine macrobenthic communities. These specimens were distributed in 3 groups in order to originate the following assembled macrobenthic communities: AMC1 was composed by 9 morphospecies of 9 specimens (one of each) (5 annelids, 3 crustaceans and 1 mollusk), AMC2 by 36 morphospecies of 36 specimens (one of each) (19 annelids, 14 mollusks and 3 crustaceans) and AMC3 by 67 specimens of 18 morphospecies (10 annelids, 5 crustaceans and 3 mollusks) (Table 1).

#### Natural macrobenthic communities (NMC)

Natural communities were sampled in four sites (NMC1, NMC2, NMC3 and NMC4) of the Sado estuary, west coast of Portugal (Fig. 6B) in May 2014. Geographical coordinates of each location were: 38.48/−8.88 for NMC1, 38.47/−8.86 for NMC2, 38.50/−8.84 for NMC3 and 38.49/−8.82 for NMC4. The macrobenthic communities of the Sado estuary have been extensively studied and the diversity of the soft bottom habitats and environmental impacts provides an appropriate test case for this study. NMC1 and NMC2 are situated in the Tróia Peninsula, near the protected area of the “Sado Estuary Nature Reserve”, and are generally less exposed to direct contamination sources of anthropogenic origin, except for ferryboat wharf located near NMC2. These Tróia Peninsula sites are more influenced by tidal hydrodynamism and have lower water residence time^56^. NMC3 and NMC4 are located on the north margin of the estuary, near the industrial zone close to the city of Setübal which harbours a number of potential sources of pollution such as factories for the production of pesticides, fertilizers and pulp mill, a thermoelectric power plant, shipyards, etc.^56^. As opposed to NMC1 And NMC2, these sites have a lower hydrodynamism, therefore facilitating the retention of contaminants and sediment’s fine particles from the upper estuary. Eleven sediment samples were collected from each site (44 samples in total) using a corer sampler (110 mm diameter, 495 mm height). One sample was used for sediment’s physico-chemical characterization and the remaining 10 samples were used for macrobenthic community assessment. The later were sieved in situ through a 0.5 mm screen in order to separate the macrobenthic invertebrates, transported refrigerated to the laboratory and stored in absolute ethanol at −20°C until processing. Five samples were then randomly chosen for morphology-based identifications and the other 5 used for metabarcoding-based identifications. Morphology-based identifications were carried out in individually separated specimens as described in the previous section. Specimens for the metabarcoding approach were processed collectively as a bulk natural community, as described further below.

### Sediment characterization

At each site three sediment’s features were determined: a) organic matter content (extrapolated from total combustible carbon, TOM): sediments were dried at 60–80°C and combusted at 500 ± 25°C for 4 h; and b) fine fraction (particle size < 63 μm): determined by sieving after treating the samples with hydrogen peroxide and disaggregation with pyrophosphate.

### DNA barcoding and HTS analyses

#### Assembled communities

Standard COI barcodes were obtained for every specimen used in the AMC study, and included in the compiled reference library. A small piece of tissue (1-2 mm) from each specimen was used for DNA extraction employing Nucleospin® Tissue kit (Macherey-Nagel Inc., Bethlehem, PA, USA) according to manufacturer’s protocols. COI was amplified using the primers LoboF1 and LoboR1 (see Table 3)^25^. PCR reactions were assembled in a 25 μL volume [2 μL DNA template, 17.5 μL molecular biology grade water, 2.5 μL 10x Invitrogen buffer, 1μL 50× MgCl2 (50 mM), 0.5 μL dNTPs mix (10 mM), 1.5 μL forward primer (10 μM), 1.5 μL reverse primer (10 μM) and 0.5 μL Invitrogen Platinum Taq polymerase (5 U/μL)]. The amplification cycle was: 95°C for 5 min; 5 cycles of 94°C for 30 s, 45°C for 1 min 30 s, 72°C for 1 min; 45 cycles of 94°C for 30 s, 54°C for 1 min 30 s, 72°C for 1 min; final extension at 72°C for 5 min. PCR products were sequenced bidirectionally using an ABI 3730XL DNA sequencer.

**Table 3.**
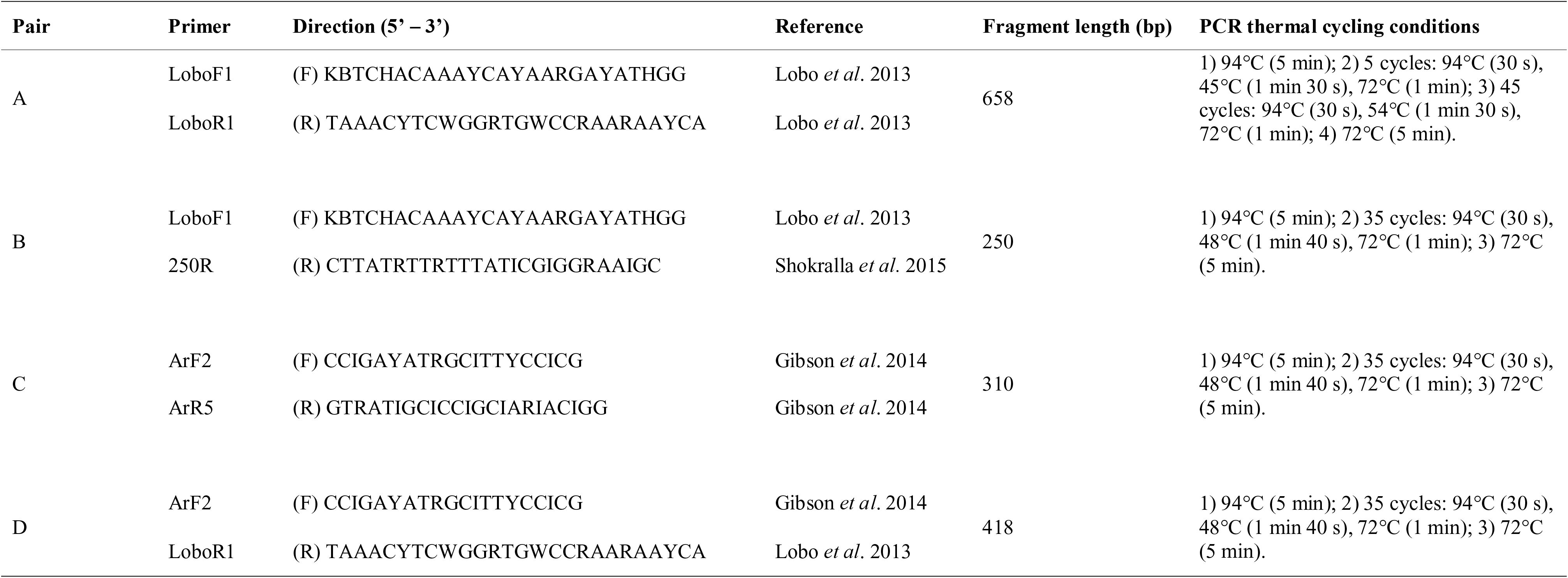
Primer pairs used to amplify COI barcode fragments from bulk samples

A reference Sanger based DNA barcode library was built using the COI sequences obtained for all 112 specimens. An in silico analyses was carried out based on Hajibabaei et al.^15^, in order to evaluate the species level discrimination ability of the various fragments sizes. Sequences from all 112 specimens were aligned using the program MEGA v.6.0^57^. Phenograms were constructed for the complete fragment (658 bp) and two fragments of 200 bp (1–200 bp and 458–658 bp) with the Neighbor-Joining (NJ) method^58^ using the Kimura 2-parameter (K2P) substitution model^59^ and 1000 bootstrap replicates. Results demonstrated that unambiguous species level identifications (intraspecific divergences below 3%) are possible even for short fragments of 200 bp (data not shown).

After tissue subsampling from individuals for building Sanger based barcoding library, the rest of the specimens were then grouped in three AMC as described above (Table 1), and each bulk sample was homogenized in 95% ethanol using a conventional blender. The homogenates were incubated at 56°C for approximately two hours to evaporate residual ethanol. Total genomic DNA of each AMC’s homogenate was extracted using Nucleospin tissue kit (Macherey-Nagel Inc.) according to manufacturer’s instructions. Four primer pairs (A, B, C and D) were used for independent amplification of either multiple fragments of CO1 barcoding region, ranging from 310 bp to 658 bp (see Table 3). PCR thermal cycling conditions for each primer pair are also presented in Table 3.

The generated amplicons from each assembled community were purified using Qiagen MiniElute PCR purification columns and eluted in 30 *μ*L molecular biology grade water. The purified amplicons from the first PCR were used as templates in the second PCR with the same amplification condition used in the first PCR with the exception of using Illumina-tailed primers in a 30-cycle amplification regime. PCR products were visualized on a 1.5% agarose gel to check the amplification success. All generated amplicons were dual indexed and pooled into a single tube and sequenced on a Miseq flowcell using a V2 Miseq sequencing kit (250 × 2) (FC-131-1002 and MS-102-2003). All PCRs were done using Eppendorf Mastercycler ep gradient S thermalcyclers and negative control reactions (no DNA template) were included in all experiments. All sequencing data generated will be deposited to Genbank and Dryad upon manuscript acceptance.

The Illumina generated reads from all COI fragments were merged with SEQPREP software (https://github.com/jstjohn/SeqPrep) requiring a minimum overlap of 25bp and no mismatches for all primer pairs, except for primer pair A (658bp fragment) resulting in paired-end reads. For primer pair A, the forward and reverse sequences were quality filtered and then concatenated in a single file before taxonomic assignment. The paired-end reads were filtered for quality using PRINSEQ software^60^ with a minimum Phred score of 20, window of 10, step of 5, and a minimum length of 100bp. USEARCH v6.0.307^61^ with the UCLUST algorithm was used to dereplicate and cluster the remaining sequences using a 99% sequence similarity cutoff. This was done to denoise any potential sequencing errors prior to further processing. Chimera filtering was performed using USEARCH with the ‘de novo UCHIME’ algorithm^62^. At each step, cluster sizes were retained, singletons and doubletons were not included for further analysis. Usable reads were compared against the reference Sanger based DNA barcode library (112 specimens) and assign to a species when displaying ≥ 98% similarity.

#### Natural communities

DNA extraction, amplification, and HTS of each natural community was carried out as described above for assembled communities. For each of the 20 bulk community samples (4 sites x 5 samples per site), two independent amplifications were performed using the primer pairs B and D. These two primers pairs were selected among the 4 previously tested in the AMC step, because the results together achieved were sufficient to obtain the maximum species recovery rates observed (see below). Amplicons obtained for each of the five samples per site were tagged separately and submitted to HTS in an Illumina MiSeq platform as described in the AMC section. After quality and size filtering, usable reads were first compared against our local barcode reference library and assign to a species when displaying ≥ 97% similarity. Reads without matching sequences in the reference library were then compared against GenBank using the same minimum threshold for taxonomic assignment. Only reads with a species match, either against the reference library or GenBank, were used in the remaining data analyses.

### Community analyses

Non-metric multidimensional scaling (nMDS) was conducted, using PAST version 3.07^63^, to show the spatial distribution of the four NMC. Bray-Curtis’s similarity index for absence-presence of species was used in order to compare morphological identification and HTS data, avoiding affecting the number of null values between samples.

Azti’s Marine Biotic Index (AMBI)^14^ is a widely used biotic index to assess the quality of benthic macroinvertebrate communities considering five ecological groups (EG) to which the benthic species are allocated. EG-I: species very sensitive to organic enrichment and present under unpolluted conditions; EG-II: species indifferent to enrichment; EG-III: species tolerant to excess organic matter enrichment; EG-IV: second-order opportunistic species; and EG-V to first-order opportunistic species (V). Because the calculation of the original AMBI index requires species abundance data, an alternative AMBI based only on presence (p) and absence (a) data (p/a AMBI) must be applied when using metabarcoding-derived species inventories, as described in Aylagas et al.^26^. The classifications obtained are somewhat similar using either p/a AMBI or the original AMBI, meaning that species relative abundance does not appear to greatly affect the outcome of the benthic assessments using this biotic index^26^. Since in our study species abundances were only available from the morphological inventories, we applied AMBI to the data from the morphology-based identification, metabarcoding and the combination of both methodologies, using the presence and absence of species to enable results’ comparison. In addition, the original AMBI index based on the abundance of specimens was also applied to the morphology-based identifications in order to validate the results.

## Acknowledgements

This work was supported by FEDER through POFC-COMPETE by national funds from ‘Fundação para a Ciência e a Tecnologia (FCT)’ in the scope of the grant FCOMP-01-0124-FEDER-015429 and also by the strategic programme UID/BIA/04050/2013 (POCI-01-0145-FEDER-007569) also funded by national funds through the FCT I.P. and by the ERDF through the COMPETE2020 - Programa Operacional Competitividade e Internacionalização (POCI). JL was supported by a PhD fellowship (SFRH/BD/69750/2010) from FCT. The authors would like to thank Stephanie Boilard (Biodiversity Institute of Ontario) for her support in the lab work.

## Author Contributions Statement

J.L. and F.O.C. wrote the manuscript. J.L., S.S., M.H. and F.O.C. globally designed the study. J.L. and M.H.C. carried out the sampling collection and specimen processing. J.L. and S.S. performed the molecular and data analyses. All authors contributed for the results’ discussion, and manuscript revision and editing.

## Supplementary information legends

**Supplementary Figure S1: Species composition of all samples of NMC.** A shows species composition considering only specimens morphologically identified to the species level. B considering specimens morphologically identified to a higher taxonomic level. C shows species composition recovered through HTS. No specimens were identified to the species level in NMC2.7. No specimens were collected in NMC2.10.

**Supplementary Table S1: Number of reads assigned to species in each NMC and primer pair.**

**Supplementary Table S2: Taxonomic classification of the species identified in each NMC through HTS (with the primer pairs B and D) and morphological identifications.** Numbers indicate the number of reads (HTS) and number of specimens for each species.

